# Boosting Motor Cortex Plasticity through Respiratory Phase-Triggered Paired Associative Stimulation

**DOI:** 10.1101/2025.01.29.635613

**Authors:** Stuart N Baker, Boubker Zaaimi

**Affiliations:** Faculty of Medical Sciences, Newcastle University, Newcastle upon Tyne NE2 4HH, UK; College of Life and Health Sciences, Aston University, Birmingham B4 7ET, UK

**Keywords:** Neural Plasticity, Paired Associative Stimulation (PAS), Motor Cortical Plasticity, Respiratory Phase Synchronization, Stroke Rehabilitation

## Abstract

Paired Associative Stimulation (PAS) has shown promise in promoting motor cortex plasticity by using transcranial magnetic stimulation (TMS) paired with peripheral nerve stimulation. However, the effectiveness of PAS is often limited by its short-lived potentiation effects. Recent research indicates that respiratory rhythms can influence cortical excitability, suggesting a potential method to enhance PAS efficacy. This study investigated the impact of synchronizing PAS with respiratory phase transitions - specifically, the transition from inspiration to expiration (I-E) and expiration to inspiration (E-I) - on motor cortical plasticity. We conducted experiments with 18 healthy volunteers (13 females, 5 males) aged 21-45 years, assessing motor-evoked potentials (MEPs) elicited by TMS applied to the left motor cortex. Participants underwent PAS sessions where paired stimuli were delivered either at I-E or E-I transitions, or at random intervals. MEPs were recorded at baseline, immediately post-PAS, and at 10, 20, and 30 minutes post-stimulation. Results showed that PAS triggered at the I-E transition significantly increased MEP amplitudes, with significant differences in MEP amplitudes at 20 minutes post-PAS between the I-E and the other conditions. This highlights the benefit of timing PAS with the I-E transition for enhanced motor cortical plasticity. These findings underscore the potential of integrating respiratory rhythms into neuromodulation techniques to improve therapeutic outcomes. Synchronizing PAS with natural respiratory phases may enhance motor recovery strategies and offers a refined approach for therapeutic interventions. This approach could be particularly relevant for stroke rehabilitation, where enhancing motor cortical plasticity is crucial for recovery.

**Significance Statement:** This study demonstrates that syncing brain stimulation with breathing patterns can enhance motor learning. By coordinating Paired Associative Stimulation (PAS)—a technique that stimulates sensory and motor inputs—with specific phases of the breathing cycle, we observed stronger responses in the motor cortex. This approach not only improves our understanding of brain adaptability but also offers a new way to fine-tune therapeutic techniques. For stroke patients, where regaining motor function is critical, integrating natural body rhythms into treatment could lead to more effective and personalized rehabilitation strategies.

## Introduction

Neural plasticity is crucial for motor recovery post-stroke (Dimyan and Cohen, 2011). Following a stroke, significant reorganization of sensorimotor cortex connections has been documented (Nudo and Milliken, 1996), and motor learning has been shown to reinforce synaptic connections in the motor cortex (Kami et al., 1995; Katai et al., 2023). Undamaged motor cortex regions undergo time-dependent plastic reorganization (Murata et al., 2015), suggesting therapeutic potential for enhancing neural connections to support improved movement recovery (Su and Xu, 2020). Most recovery occurs within the first three months when plasticity is heightened, yet many trials on plasticity protocols are conducted during the chronic stage post-stroke (Krakauer and Carmichael, 2017). Various brain stimulation techniques have shown promise in promoting plasticity in chronic stroke patients (Elsner et al., 2016; Fridriksson et al., 2018) though the effects reported have been moderate.

Paired associative stimulation (PAS) has shown success in inducing plastic changes in the motor cortex of both healthy and chronic stroke patients (Silverstein et al., 2019). Stefan et al. (Stefan et al., 2000) introduced PAS, where transcranial magnetic stimulation (TMS) of the motor cortex is consistently preceded by stimulation of the contralateral median nerve. An increase in motor-evoked potentials (MEPs) after PAS indicates heightened excitability in the stimulated motor cortex, suggesting LTP-like mechanisms mediated by NMDA receptors (Stefan et al., 2002; Volz et al., 2016). However, the LTP-like effect typically lasts only around 90 minutes (Michael et al., 2007), insufficient for sustained therapeutic benefit. Efforts to enhance PAS’s effect on plasticity have explored modulation by serotonin (Batsikadze et al., 2013) and network excitability manipulated by transcranial direct current stimulation (Michael et al., 2007).

Recently a broad bottom-up influence of the cortical excitability by the breathing rhythm has been demonstrated in memory (Schreiner et al., 2023), and attention (Zelano et al., 2016; Kluger et al., 2021). Each breath appears to bring a tidal wave of excitability to cortical neurons (Ito et al., 2014). It has been demonstrated that breathing drives oscillations of the membrane potential of neurons in various cortical areas including prefrontal and somatosensory cortices (Juventin et al., 2023). A study (Li and Rymer, 2011) examined the relationship between respiration and motor activity by using TMS or electrical stimulation on hand muscles and discovered a general enhancement of the motor system related to breathing. The authors observed that delivering stimulation to finger extensors during voluntary inspiration resulted in a significant decrease in finger flexor spasticity in a stroke patient. Furthermore, another study (Kluger and Gross, 2020) has demonstrated higher beta cortico-muscular coherence during natural breathing compared to voluntary deep breathing, and that this coherence exhibits cyclic modulation depending on the phase of respiration.

The transition from inspiration to expiration and vice versa is critical because these shifts in the body’s physiological state can modulate cortical excitability. A study by Zelano et al. (Zelano et al., 2016) demonstrated that the act of inhaling through the nose significantly increases delta oscillations and enhances cognitive functions such as memory recall, directly linking respiratory phase to cortical activity. Conversely, Nakamura et al. (Nakamura et al., 2018) suggest that the transition from expiration to inspiration might affect reaction time and recall accuracy in a delayed match-to-sample visual recognition task. Their results showed that when this transition occurred between image presentation and response, subjects experienced significantly longer response times and decreased recall accuracy compared to trials where this transition did not occur.

The transitions between the breathing phases are particularly relevant for neuromodulation techniques like PAS. By timing PAS protocols to align with the transition from inspiration to expiration or vice versa, it is possible to exploit these natural physiological states to enhance or decrease cortical excitability, respectively. We investigated breathing’s role in modulating PAS-induced LTP-like effects. This study assesses the impact of timing PAS with breathing phases on motor cortical plasticity, focusing on transitions between inspiration and expiration.

## Methods

### Participants

The study included 18 healthy volunteers (13 females, 5 males; aged 21–45 years), each participating in 3 sessions with at least 24 hours between sessions, totalling 54 sessions. All procedures were approved by the Newcastle University Medical School’s ethical committee. Written informed consent was obtained from all participants after detailed explanation of the procedures. Participants were seated comfortably with their right forearm on an adjacent table. They were instructed to remain relaxed, breathe through their nose, and watch a documentary, with EMG and respiration monitored throughout.

### EMG Recording

EMG recordings were obtained from the right abductor pollicis brevis (APB) using adhesive surface electrodes (H59P Kendall; Covidien, Dublin, Ireland) placed over the muscle belly (1 cm apart). Electrodes were connected to a D360 amplifier (Digitimer Ltd, Welwyn Garden City, UK) with a gain of 1000 and a bandpass filter from 30 Hz to 2 kHz. The signal was digitized via a micro1401 laboratory interface (Cambridge Electronics Design, Cambridge, UK) and stored using Spike2 software. The EMG signal was rectified prior to analysis.

### Median Nerve Stimulation

The right median nerve was stimulated through bipolar electrodes at the wrist (cathode proximal) using a 0.5 ms square wave pulse. Stimulation intensity was set at three times the perceptual threshold with a DS7A constant-current isolated stimulator (Digitimer Ltd, Welwyn Garden City, UK).

### Respiration

Participants wore surgical masks with a temperature sensor (LM34, Texas Instruments) positioned below the nostrils. The mask was adjusted for comfort to prevent direct skin contact. The sensor signal was powered, amplified with a custom circuit, and digitized via a micro1401 laboratory interface (Cambridge Electronics Design, Cambridge, UK). A custom Spike2 script enabled online detection of respiratory phase transitions (I-E or E-I) to trigger paired stimulations. In control sessions, stimulations were triggered at random intervals between 9.4 and 12.5 seconds.

Figure 1 illustrates respiratory signals collected during the study, including stimulation timings and alignment with respiratory transitions. Figure 1A and Figure 1E show paired stimulations triggered at the I-E and E-I transitions, respectively. Figure 1B and Figure 1F display respiratory traces aligned with these transitions. Figure 2C and Figure 2D present data from sessions with random stimulation intervals, with the respiratory cycles aligned to indicate varied trigger points.

**Figure 1.**
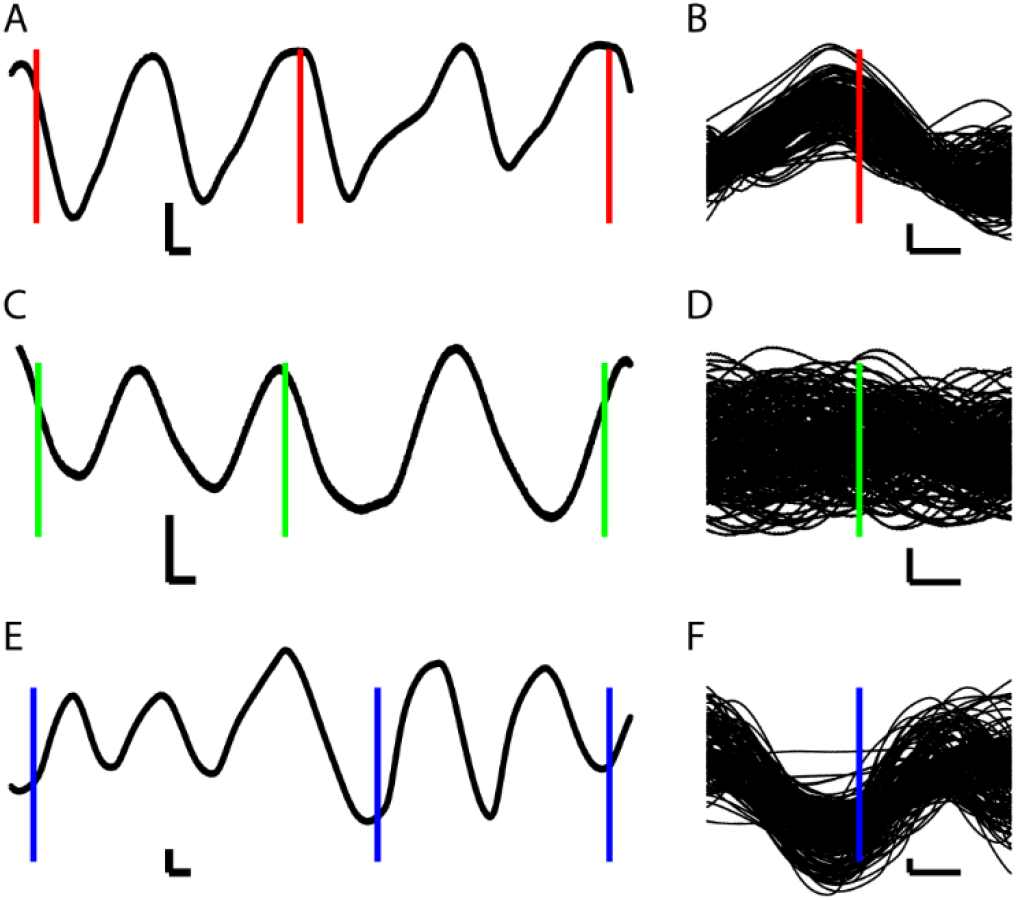
Triggering PAS with Respiratory Phases from One Participant. Panels A, C, and D display the raw respiratory signals measured by the thermoresistor, along with the timing of three consecutive paired stimulations triggered at the I-E transition (A), randomly (C), and at the E-I transition (D). The voltage increases during inspiration and decreases during expiration. Panels B, D, and F show the overlay of respiratory traces aligned with the paired stimulation (TMS) triggered at the I-E transition (B), random intervals (D), and the E-I transition (F). Calibration bars: horizontal = 1 s, vertical = 100 mV.

**Figure 2.**
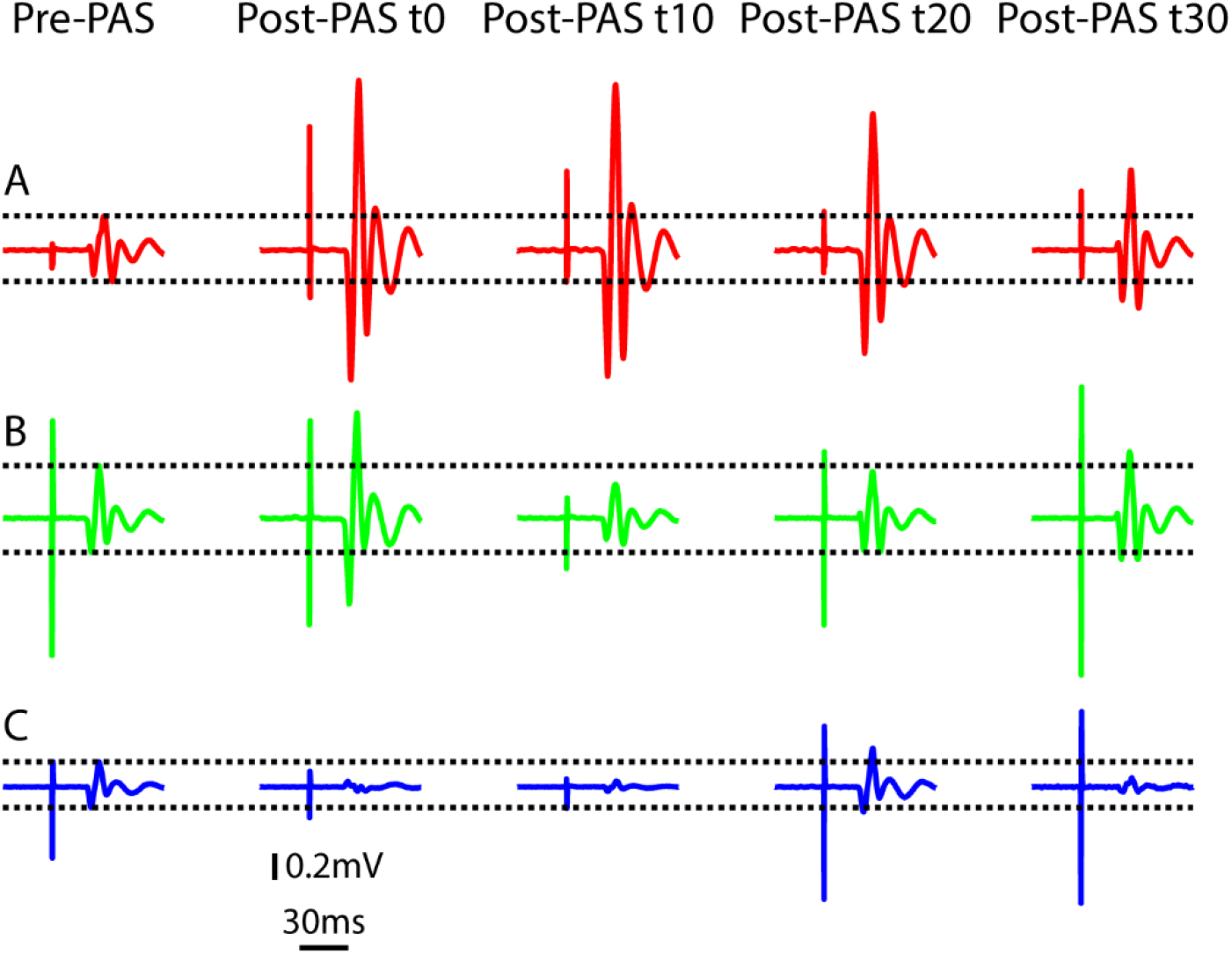
MEP Traces from One Participant. Panel A shows MEPs with paired stimuli at the I-E transition, demonstrating increased amplitude that remains elevated for 30 minutes. Panel B depicts MEPs with random stimuli, showing a brief increase followed by a return to baseline. Panel C illustrates decreased MEP amplitude with stimuli triggered at the E-I transition. Each panel includes averages from 20 TMS stimulations pre-PAS, immediately post-PAS, and at 10, 20, and 30 minutes post-PAS.

### TMS Stimulation

TMS was applied using a round coil connected to a Magstim 2002 stimulator (Magstim Ltd, Whitland, UK) centered over the vertex to stimulate the left motor cortex and elicit MEPs in the right APB. The resting motor threshold (RMT) was determined as the minimum stimulator output inducing MEPs of more than 0.1 mV peak-to-peak in at least 5 of 10 single shock stimulations. Throughout the recording, TMS was delivered at 1.2 x RMT. Baseline MEPs were recorded with 20 single pulse TMS at 5-second intervals (Pre-PAS). Following this, 200 paired stimulations were administered: median nerve stimulation followed by single pulse TMS after a 25-millisecond delay. Each session’s paired stimulations were either triggered at I-E or E-I transitions or randomly (control session). After the 200 paired stimulations, 20 MEPs were recorded at t0 (immediately post-PAS), and additional sets were recorded at t10, t20, and t30 minutes post-stimulation. Sessions were conducted in a pseudo-random order.

### Statistical analysis

Data were analyzed using a combination of t-tests, Fisher’s exact tests, and ANOVA to assess the effects of post-PAS time, respiratory phase, and their interactions on MEP amplitudes. A repeated-measures ANOVA was employed to evaluate the main effects and interactions of time (pre-PAS, t0, t10, t20, t30) and respiratory phase (I-E, E-I, control) on MEP sizes. This test determined whether there were statistically significant differences in MEP amplitudes across different post-PAS time points and between respiratory phase conditions. Paired t-tests were conducted to assess whether post-PAS MEP amplitudes significantly differed from pre-PAS baseline levels at each post-PAS time point (t0, t10, t20, t30) for each respiratory phase condition. The t-test was also used to compare the MEPs between conditions at specific time periods. Fisher’s exact test was applied to assess the significance of categorical data, such as the distribution of responders versus non-responders across different conditions, which was particularly useful for small sample sizes and provided insights into the associations between respiratory phase transitions and the likelihood of MEP facilitation. To ensure the validity of parametric tests, the normality of the data was assessed using a Q-Q plot and the Kolmogorov-Smirnov test in MATLAB. The p-value from the Kolmogorov-Smirnov test was greater than 0.05, indicating no significant deviation from normality. Additionally, the Q-Q plot showed that the data points closely followed the reference line, further supporting the assumption that the data is normally distributed.

## Results

Figure 2 presents the raw MEP traces from the APB muscle evoked by TMS in one participant. The most significant increase in MEP amplitude is observed when paired stimuli are triggered at the I-E transition (Fig. 2A). This increase persists above the average pre-PAS MEP levels for up to 30 minutes following the pairing. In contrast, when paired stimuli are triggered randomly throughout the respiratory cycle, as in conventional PAS experiments (Fig. 2B), MEPs show a transient increase immediately after pairing, followed by a return to pre-PAS amplitude levels. Notably, when paired stimuli are triggered at the E-I transition, as shown in Figure 2C, there is a decrease in MEP amplitude following the PAS protocol.

Figure 3 presents MEP data from individual participants recorded at four post-pairing intervals: immediately after, 10 minutes, 20 minutes, and 30 minutes. MEP amplitude is represented as a percentage, with values exceeding 100% indicating facilitation. The data reveal significant variability among participants, with some demonstrating facilitation and others exhibiting suppression effects.

**Figure 3.**
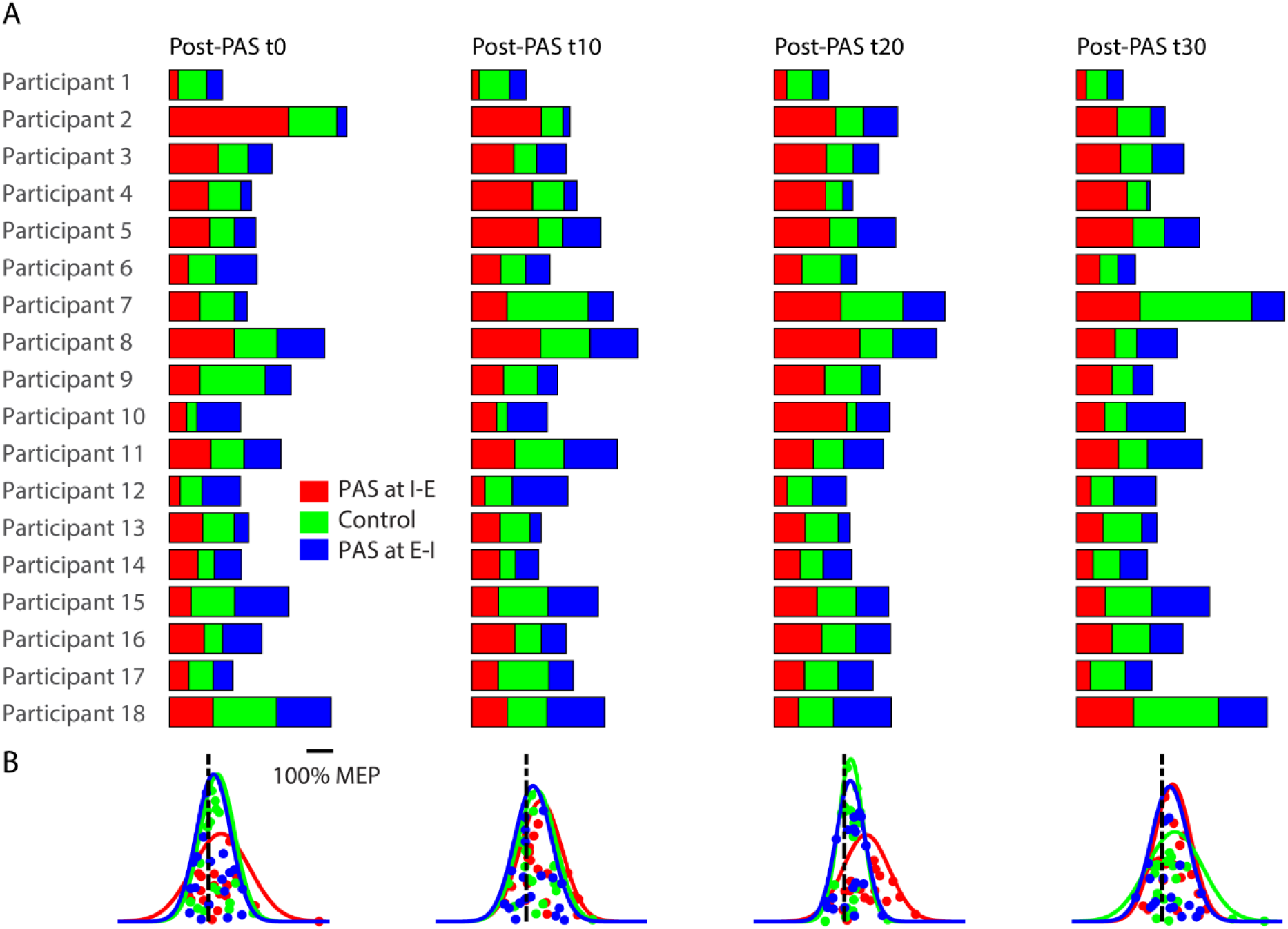
Distribution of MEP Amplitudes Across Protocols and Time Points. Distribution of Motor Evoked Potential (MEP) amplitudes for individual participants across three experimental conditions (PAS at I-E, Control, PAS at E-I) at four post-pairing time points: t0 (immediately after pairing), t10 (10 minutes post-pairing), t20 (20 minutes post-pairing), and t30 (30 minutes post-pairing). MEP amplitudes are expressed as a percentage relative to the pre-pairing baseline (100%), with values above 100% indicating facilitation. Panel A: Bar plots depict the average MEP amplitude for each protocol at each time point. Panel B: Normal distributions are plotted to show the spread of MEP amplitudes within each protocol condition. At t20, the I-E transition condition shows a distinct peak, indicating a pronounced facilitation effect. By t30, the distributions for all protocols converge, reflecting similar MEP amplitudes across conditions.

For instance, at Time Point t0 (Immediately After Pairing), Participant 5 showed a marked facilitation under the I-E protocol compared to the Control (160% ± 15% vs 97% ± 8%), with a t-statistic of 2.25 and p = 0.04, indicating a strong effect of the I-E transition. Conversely, Participant 7 did not exhibit significant differences between the I-E and Control conditions (119% ± 14% vs 136% ± 20%), with a t-statistic of -0.9 and p = 0.38. However, Participant 7 did show a significant increase when comparing the Control to E-I (136% ± 20% vs 50% ± 5%), with a t-statistic of 2.7 and p = 0.01. These findings demonstrate variability in responses, with some participants showing clear facilitation under specific protocols while others did not.

At Time Point t10 (10 Minutes Post-Pairing), the t-test results indicated that Participant 3 experienced a significant increase in MEP amplitude under the I-E protocol compared to the Control (165% ± 11% vs 90% ± 8%), with a t-statistic of 2.93 and p = 0.01, reflecting sustained effects of the I-E transition. In contrast, Participant 12 did not show significant differences between conditions (e.g., I-E vs. Control: 51% ± 8% vs 107% ± 15%), with a t-statistic of -1.49 and p = 0.15, suggesting a more uniform response across protocols at this time point.

At Time Point t20 (20 Minutes Post-Pairing), Participant 8 exhibited a significant increase in MEP amplitude with the I-E protocol compared to both the Control (336% ± 61% vs 128% ± 36%), with a t-statistic of 1.39 and p = 0.18, and E-I (172% ± 20%), with a t-statistic of 1.69 and p = 0.11. This suggests that the I-E transition had a pronounced effect at this time point. Conversely, Participant 10 showed no significant difference between the I-E and Control conditions (284% ± 20% vs 37% ± 3%), with a t-statistic of 5.81 and p < 0.01 but exhibited significant changes when comparing Control to E-I (131% ± 10%), with a t-statistic of -5.44 and p < 0.01.

At Time Point t30 (30 Minutes Post-Pairing), Participant 2 demonstrated a significant increase in MEP amplitude under the Control condition compared to both I-E (132% ± 14% vs 160% ± 15%), with a t-statistic of -0.26 and p = 0.8, and E-I (55% ± 11%), with a t-statistic of 2.12 and p = 0.05, indicating a shift in response effectiveness over time. Conversely, Participant 15 showed no significant differences between conditions (e.g., Control vs. I-E: 183% ± 14% vs 112% ± 9%), with a t-statistic of -3.18 and p = 0.01, suggesting that the effects of the different protocols were less pronounced or more consistent for this participant at the later time point.

Figure 4 illustrates the mean MEP amplitudes relative to pre-PAS values across three experimental conditions (I-E, Control, E-I) measured at four distinct time points (0, 10, 20, and 30 minutes post-pairing). Error bars represent the standard error of the mean (SEM), depicting the variation in MEP amplitudes for each condition over time. At the 20-minute time point (t20), significant differences were observed using t-tests. Specifically, there was a significant difference between the I-E and Control conditions, with a t-statistic of 2.43 and a p-value of 0.0263. Additionally, a significant difference was found between the I-E and E-I conditions, with a t-statistic of 2.65 and a p-value of 0.0167. These results suggest that at this time point, the I-E condition has a significantly different impact on MEP amplitudes compared to both the Control and E-I conditions. Overall, while the experimental conditions show a significant effect on MEP amplitudes, with notable differences at t20, the effect of time across all conditions is not significant, as indicated by an F-value of 0.13 and a p-value of 0.9402. Moreover, there is no significant interaction between conditions and time points, with an F-value of 0.57 and a p-value of 0.7511. This suggests that although different conditions influence MEP amplitudes (F-value of 3.05 and a p-value of 0.0495), the temporal changes in MEP responses do not differ significantly across conditions.

**Figure 4.**
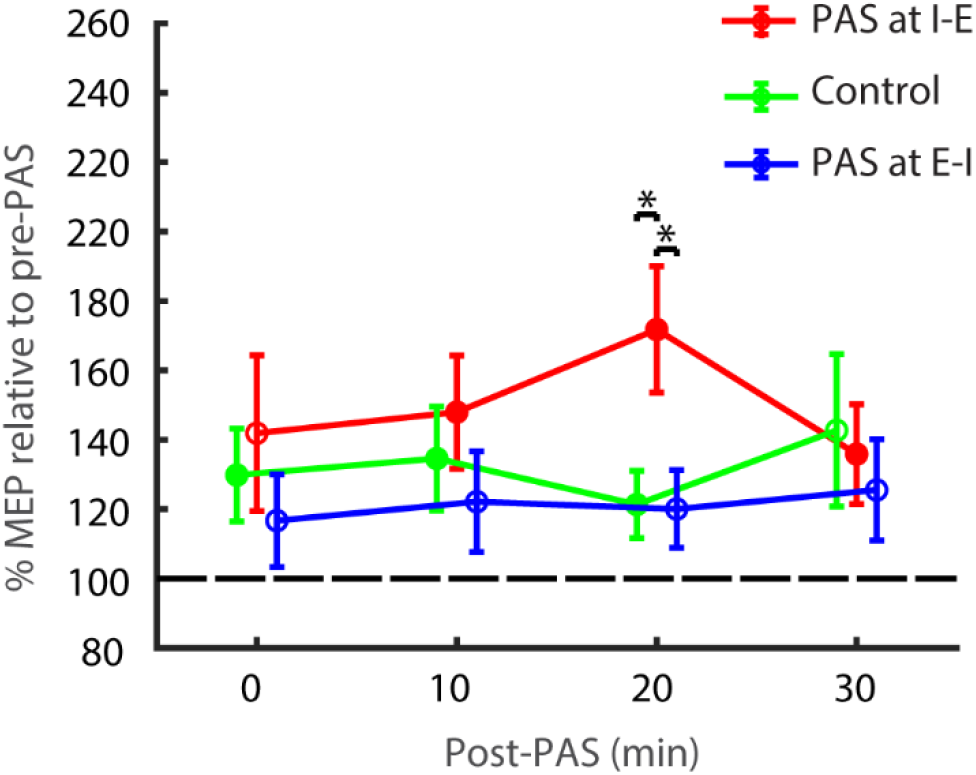
Mean MEP amplitudes relative to pre-PAS values for three conditions (I-E, Control, E-I) at four time points (0, 10, 20, 30 minutes post-pairing). Error bars denote SEM. Stars indicate significant differences (t test, p < 0.05). ANOVA analysis showed that the conditions significantly affect MEP amplitudes, but the effect of time and the interaction between conditions and time points are not significant.

Figure 5 depicts the categorization of participants into responders and non-responders at each time point based on the significance of their Motor-Evoked Potential (MEP) amplitudes relative to pre-Pulse After Stimulation (PAS) values. Responders are defined as those showing significant changes (p < 0.05), while non-responders show no significant change (p > 0.05). The figure illustrates variations in the proportions of responders versus non-responders across different conditions. It specifically highlights whether participants who did not respond to the Control condition at a given time point subsequently showed significant responses under the PAS conditions of I-E or E-I. This comparison sheds light on how aligning the pairing trigger with I-E or E-I conditions affects the responsiveness of participants who initially did not respond to the Control.

**Figure 5:**
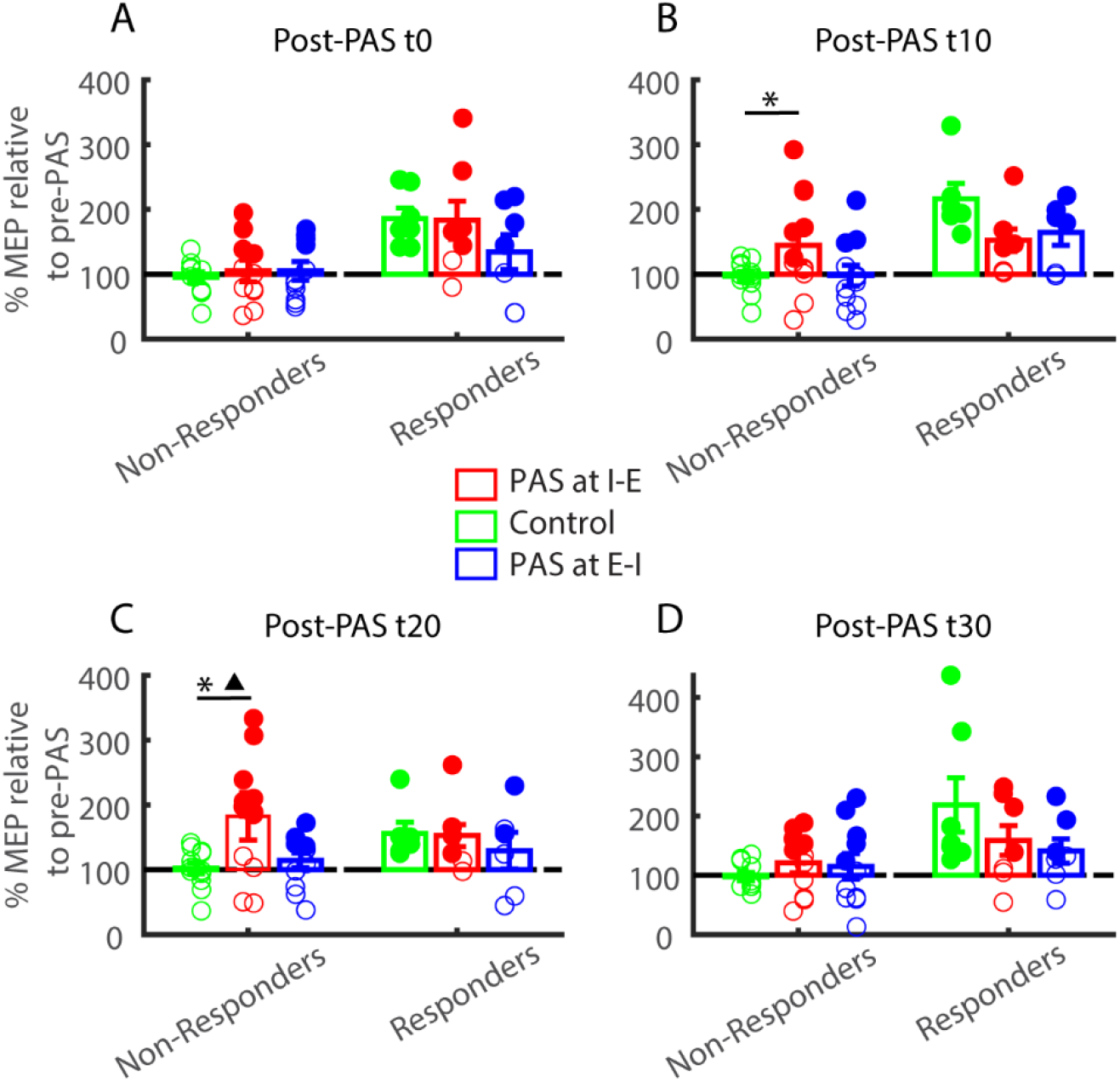
Mean relative post-PAS Motor-Evoked Potential (MEP) amplitudes expressed as percentages at four time points (0, 10, 20, and 30 minutes post-pairing). Bar plots depict the mean MEP amplitudes for three conditions: Control, I-E, and E-I, with error bars representing the standard error of the mean. Individual data points are jittered for clarity and color-coded based on significance, with filled markers denoting significant deviations from pre-PAS values (p < 0.05) and open markers indicating non-significant changes (p > 0.05). At t20, the mean MEP amplitude was significantly higher in the I-E condition compared to the Control, as marked by the triangle. Additionally, the proportion of responders increased significantly in the I-E condition compared to the Control at t10 and t20 (denoted by a triangle p<0.01).

To assess the statistical significance of the increase in responders under the I-E or E-I conditions compared to the Control condition, Fisher’s Exact Test was performed. The results showed a statistically significant increase in the proportion of responders (denoted by a star, Fisher’s Exact Test, p < 0.01) at time points t10 and t20 when the paired stimulation was aligned with the I-E transition, compared to the Control condition.

Additionally, for non-responders at t20, the mean response was significantly higher under the I-E condition compared to the Control condition, as indicated by a triangle (average ± SEM: Control vs. I-E, 102.41% ± 8.64% vs. 182.44% ± 25.86%, t-statistic = -2.86, p = 0.0154). This result demonstrates a notable increase in the mean response for non-responders at t20 when tested with the I-E pairing transition.

## Discussion

### Enhancing Motor Cortical Plasticity with Respiratory Phase Synchronization

This study provides significant insights into how synchronizing Paired Associative Stimulation (PAS) with respiratory phase transitions, particularly from inspiration to expiration (I-E), can significantly enhance motor cortical plasticity. Our results reveal that aligning PAS with the I-E transition phase markedly improves motor-evoked potentials (MEPs), with the most pronounced effects observed 20 minutes post-stimulation. This finding underscores the critical role of respiratory rhythms in modulating cortical excitability and suggests a promising strategy for refining neuromodulation therapies, particularly in stroke rehabilitation.

### Breathing and Motor Control: Complex Interactions

The synchronization of motor activities with breathing rhythms has been documented in various studies. For instance, research demonstrated that both lower limb locomotion and upper limb actions, such as finger tapping, exhibit a rhythmic coordination with breathing patterns (Wilke et al., 1975; Siegmund et al., 1999). This synchronization is not merely coincidental but indicates an underlying neural mechanism that integrates respiratory rhythms with motor control.

A recent study has further elucidated this relationship showing that voluntary movements are more likely to occur during expiration (Park et al., 2020). Ravignani and Kotz have expanded this understanding by showing that rhythmic vocalizations and movements are interconnected, suggesting that rhythmic breathing may influence not only vocal but also motor patterns (Ravignani and Kotz, 2020). This is particularly relevant in the context of understanding how respiratory rhythms shape both the initiation and execution of voluntary movements.

### Integration of Breathing with Motor Control

The integration of breathing with motor control involves complex neural pathways, influenced by distinct respiratory phases and their associated neural mechanisms. The respiratory cycle consists of two main phases: inspiration and expiration. These phases are controlled by separate neural populations in the brainstem. The onset of inspiration, marked by the transition from expiration to inspiration (EI), is driven by the PreBötzinger complex (PreBötC) in the medulla oblongata, which acts as the primary inspiratory rhythm generator (Smith et al., 2013). In contrast, the transition from inspiration to expiration (IE) involves a more gradual onset, regulated by mechanisms such as the “inspiratory off-switch” in the pons and the modulation of neural excitability (Dutschmann and Dick, 2012; Richter and Smith, 2014).

Efferent projections from brainstem respiratory centers, including the periaqueductal grey nucleus and the preBötzinger complex, influence various brain regions such as the thalamus, basolateral amygdala, nucleus accumbens, and sensorimotor cortices (Koutsikou et al., 2015; Yang and Feldman, 2018). These networks enable continuous updates on breathing states and integrate them with sensorimotor outputs. While the influence of breathing on motor control is well-documented in the contexts of martial arts and physical training (Gariépy et al., 2010; Boyadzhieva and Kayhan, 2021), its effects on movement execution and plasticity modulation remain underexplored.

Recent findings by Nakamura (Nakamura et al., 2022) provide valuable insights into how these distinct respiratory transitions affect brain activity. Nakamura’s study observed differential effects of these transitions on brain regions involved in attention and memory. Specifically, during the EI transition, there was reduced activation in areas such as the right temporoparietal junction, as well as in the left and right middle frontal gyrus, dorsomedial prefrontal cortex, and somatosensory areas, which are integral to attentional control and memory processing. This pattern suggests that the EI transition might interfere with cognitive functions related to information manipulation.

Interestingly, our research aligns with these findings by demonstrating that triggering PAS at the I-E transition—when the brain’s neural activity shifts from inspiration to expiration— enhances motor cortex plasticity. This timing appears to optimize the stimulation’s impact, suggesting that the I-E transition, marked by a gradual and controlled neural shift, is particularly conducive to modulating motor cortex responses. The alignment of PAS with the I-E transition offers a promising approach to leveraging the body’s natural rhythms to enhance motor learning and plasticity.

### Variability in PAS and Implications for Clinical Application

Despite its potential, PAS faces significant challenges in achieving consistent clinical application. Variability in both animal models and human studies has limited PAS’s widespread adoption. For instance, a study using a freely behaving rat model tested various interstimulus intervals with the expectation that they would induce corticomotor potentiation based on spike timing-dependent plasticity (STDP) principles. However, despite numerous stimulus pairings, the study failed to demonstrate reliable changes in corticospinal excitability compared to control conditions, questioning the effectiveness of PAS under these experimental conditions (Ting et al., 2020).

One major issue is that PAS does not always translate effectively to in vivo conditions due to the complexity of neural circuits and ongoing spontaneous and behavior-related activity (Markram et al., 1997). This complexity introduces variability that can disrupt the precise timing required for STDP, complicating the application and effectiveness of PAS. Additionally, high intersubject variability in PAS responses observed in clinical trials (McGie et al., 2014; Tarri et al., 2018) suggests that PAS effects are not consistently reproducible, further complicating its practical use. Sale et al. (Sale et al., 2007) demonstrated that PAS was more effective in the afternoon compared to the morning, potentially due to fluctuations in cortisol levels affecting neuroplasticity. This finding highlights the need for standardized experimental conditions to reduce variability and improve reproducibility. Considering these challenges, our study offers a potential pathway to enhance PAS efficacy. By aligning PAS with the I-E transition, we have demonstrated that it is possible to make non-responders to traditional PAS protocols respond more reliably. This alignment leverages the rhythmic physiological changes in breathing patterns to synchronize stimulation with key respiratory phases, potentially enhancing the effectiveness of PAS.

### Timing and Mechanisms of PAS Effects

Our study highlights that PAS has the most pronounced effects 20 minutes after stimulation, particularly when timed with the I-E transition phase rather than the E-I transition. Although traditional PAS triggers an immediate facilitation response, synchronizing PAS with the end of the inspiratory phase induces neuromodulatory effects, supporting the development of long- term potentiation (LTP) that becomes evident at the 20-minute mark. The observed boost in facilitation 20 minutes post-pairing is likely due to the gradual progression of synaptic changes and intracellular signaling pathways related to LTP, which are optimized by the transition from inspiratory to expiratory phases.

Breathing significantly influences brain activity, affecting neural oscillations and motor actions. In rodents, breathing rhythms synchronize with brain oscillations across delta, theta, and gamma frequencies in various brain regions (Heck et al., 2019). Human studies also reveal respiration-coupled neural activity, particularly in the piriform cortex and hippocampus (Herrero et al., 2018). Respiratory coupling is linked to cognitive performance through theta-gamma interactions (Köster et al., 2014; Schomburg et al., 2014; Tamura et al., 2017), and specific phases of exhalation enhance performance in tasks like finger manipulation (Nakamura et al., 2018). Aligning PAS with respiratory rhythms, particularly the I-E transition phase, may optimize neuroplastic effects and illustrate the importance of timing PAS to enhance therapeutic outcomes.

### PAS and Breathing in Stroke Rehabilitation

PAS represents a promising approach in stroke rehabilitation, leveraging non-invasive brain stimulation to modulate cortical excitability and promote neuroplasticity. The effectiveness of PAS, especially when combined with repeated sessions and rehabilitative treatments, remains a subject of ongoing research. Single-session studies have shown increased motor cortex excitability and improved motor performance (Wessel et al., 2015). However, the benefits of prolonged or combined treatments are less certain. For example, Sui et al. (Sui et al., 2021) reported significant improvements in subacute stroke patients using PAS, suggesting potential benefits for motor function. In another study, over four weeks of paired stimulation, nine patients with chronic stable hemiparesis showed improvements in neurophysiological and functional measures, indicating potential therapeutic benefits (Uy et al., 2003). However, further research is needed to determine optimal PAS protocols and patient profiles.

On the other hand, a recent study explored a 2-week adapted mindfulness meditation protocol for chronic stroke survivors with spasticity. Participants reported significant improvements in spasticity following the intervention, which involved controlled breathing (Wathugala et al., 2019). Integrating respiratory phase considerations into rehabilitation protocols could further enhance PAS effectiveness by aligning with natural physiological rhythms, potentially improving motor recovery and neuroplasticity.

### Conclusion and Future Directions

In summary, aligning PAS with specific respiratory phase transitions, particularly the I-E phase, significantly enhances motor cortical plasticity. These findings open new avenues for developing advanced neuromodulation techniques and emphasize the importance of incorporating natural physiological rhythms into therapeutic interventions. Future research should focus on exploring these techniques in stroke patients, refining timing protocols, and elucidating the mechanisms underlying respiratory phase modulation to maximize therapeutic benefits. By integrating breathing rhythms into neuromodulation therapies, we may improve motor recovery and cognitive function, offering a more effective approach to rehabilitation.

## Acknowledgement

We would like to extend our sincere thanks to all the participants for their time and valuable contributions to this study. We wish to acknowledge Natalie Maffit, Tim- Joshua Andres, Maria Germann, Demetris Soteropoulos, and Terri Jackson for their essential assistance in organizing visits, coordinating the experiments. Your contributions were crucial to the success of this project.

## References

Batsikadze G, Paulus W, Kuo M-F, Nitsche MA (2013) Effect of Serotonin on Paired Associative Stimulation-Induced Plasticity in the Human Motor Cortex. Neuropsychopharmacology 38:2260–2267.

Boyadzhieva A, Kayhan E (2021) Keeping the Breath in Mind: Respiration, Neural Oscillations, and the Free Energy Principle. Frontiers in neuroscience 15:647579.

Dimyan MA, Cohen LG (2011) Neuroplasticity in the context of motor rehabilitation after stroke. Nature Reviews Neurology 7:76–85.

Dutschmann M, Dick TE (2012) Pontine mechanisms of respiratory control. Comprehensive Physiology 2:2443–2469.

Elsner B, Kugler J, Pohl M, Mehrholz J (2016) Transcranial direct current stimulation (tDCS) for improving activities of daily living, and physical and cognitive functioning, in people after stroke. Cochrane Database of Systematic Reviews.

Fridriksson J, Rorden C, Elm J, Sen S, George MS, Bonilha L (2018) Transcranial Direct Current Stimulation vs Sham Stimulation to Treat Aphasia After Stroke: A Randomized Clinical Trial. JAMA Neurology 75:1470–1476.

Gariépy J-F, Missaghi K, Dubuc R (2010) Chapter 12 - The interactions between locomotion and respiration. In: Progress in Brain Research (Gossard J-P, Dubuc R, Kolta A, eds), pp 173–188: Elsevier.

Heck DH, Kozma R, Kay LM (2019) The rhythm of memory: how breathing shapes memory function. J Neurophysiol 122:563–571.

Herrero JL, Khuvis S, Yeagle E, Cerf M, Mehta AD (2018) Breathing above the brain stem: volitional control and attentional modulation in humans. 119:145–159.

Ito J, Roy S, Liu Y, Cao Y, Fletcher M, Lu L, Boughter JD, Grün S, Heck DH (2014) Whisker barrel cortex delta oscillations and gamma power in the awake mouse are linked to respiration. Nature Communications 5:3572.

Juventin M, Zbili M, Fourcaud-Trocmé N, Garcia S, Buonviso N, Amat C (2023) Respiratory rhythm modulates membrane potential and spiking of nonolfactory neurons. 130:1552–1566.

Kami A, Meyer G, Jezzard P, Adams MM, Turner R, Ungerleider LG (1995) Functional MRI evidence for adult motor cortex plasticity during motor skill learning. Nature 377:155–158.

Katai S, Maeda M, Katsuyama S, Maruyama Y, Midorikawa M, Okushima T, Yoshida K (2023) Cortical reorganization correlates with motor recovery after low-frequency repetitive transcranial magnetic stimulation combined with occupational therapy in chronic subcortical stroke patients. Neuroimage: Reports 3:100156.

Kluger DS, Gross J (2020) Depth and phase of respiration modulate cortico-muscular communication. NeuroImage 222:117272.

Kluger DS, Balestrieri E, Busch NA, Gross J (2021) Respiration aligns perception with neural excitability. eLife 10.

Köster M, Friese U, Schöne B, Trujillo-Barreto N, Gruber T (2014) Theta–gamma coupling during episodic retrieval in the human EEG. Brain Research 1577:57–68.

Koutsikou S, Watson TC, Crook JJ, Leith JL, Lawrenson CL, Apps R, Lumb BM (2015) The Periaqueductal Gray Orchestrates Sensory and Motor Circuits at Multiple Levels of the Neuraxis. The Journal of neuroscience : the official journal of the Society for Neuroscience 35:14132–14147.

Krakauer JW, Carmichael ST (2017) Broken Movement: The Neurobiology of Motor Recovery after Stroke. In: The MIT Press.

Li S, Rymer WZ (2011) Voluntary Breathing Influences Corticospinal Excitability of Nonrespiratory Finger Muscles. 105:512–521.

Markram H, Lübke J, Frotscher M, Sakmann B (1997) Regulation of synaptic efficacy by coincidence of postsynaptic APs and EPSPs. Science (New York, NY) 275:213–215.

McGie SC, Masani K, Popovic MR (2014) Failure of spinal paired associative stimulation to induce neuroplasticity in the human corticospinal tract. The journal of spinal cord medicine 37:565–574.

Michael AN, Amelie R, Min-Fang K, Anja KF, David L, Nicolas L, Frithjof T, Walter P (2007) Timing-Dependent Modulation of Associative Plasticity by General Network Excitability in the Human Motor Cortex. The Journal of Neuroscience 27:3807.

Murata Y, Higo N, Hayashi T, Nishimura Y, Sugiyama Y, Oishi T, Tsukada H, Isa T, Onoe H (2015) Temporal Plasticity Involved in Recovery from Manual Dexterity Deficit after Motor Cortex Lesion in Macaque Monkeys. 35:84–95.

Nakamura NH, Fukunaga M, Oku Y (2018) Respiratory modulation of cognitive performance during the retrieval process. PloS one 13:e0204021.

Nakamura NH, Fukunaga M, Yamamoto T, Sadato N, Oku Y (2022) Respiration-timing-dependent changes in activation of neural substrates during cognitive processes. Cerebral cortex communications 3:tgac038.

Nudo RJ, Milliken GW (1996) Reorganization of movement representations in primary motor cortex following focal ischemic infarcts in adult squirrel monkeys. Journal of Neurophysiology 75:2144–2149.

Park H-D, Barnoud C, Trang H, Kannape OA, Schaller K, Blanke O (2020) Breathing is coupled with voluntary action and the cortical readiness potential. Nature Communications 11:289.

Ravignani A, Kotz SA (2020) Breathing, voice, and synchronized movement. Proceedings of the National Academy of Sciences of the United States of America 117:23223–23224.

Richter DW, Smith JC (2014) Respiratory rhythm generation in vivo. Physiology (Bethesda, Md) 29:58–71.

Sale MV, Ridding MC, Nordstrom MA (2007) Factors influencing the magnitude and reproducibility of corticomotor excitability changes induced by paired associative stimulation. Exp Brain Res 181:615–626.

Schomburg EW, Fernández-Ruiz A, Mizuseki K, Berényi A, Anastassiou CA, Koch C, Buzsáki G (2014) Theta phase segregation of input-specific gamma patterns in entorhinal-hippocampal networks. Neuron 84:470–485.

Schreiner T, Petzka M, Staudigl T, Staresina BP (2023) Respiration modulates sleep oscillations and memory reactivation in humans. Nature Communications 14:8351.

Siegmund GP, Edwards MR, Moore KS, Tiessen DA, Sanderson DJ, McKenzie DC (1999) Ventilation and locomotion coupling in varsity male rowers. Journal of applied physiology (Bethesda, Md : 1985) 87:233–242.

Silverstein J, Cortes M, Tsagaris KZ, Climent A, Gerber LM, Oromendia C, Fonzetti P, Ratan RR, Kitago T, Iacoboni M, Wu A, Dobkin B, Edwards DJ (2019) Paired Associative Stimulation as a Tool to Assess Plasticity Enhancers in Chronic Stroke. 13.

Smith JC, Abdala AP, Borgmann A, Rybak IA, Paton JF (2013) Brainstem respiratory networks: building blocks and microcircuits. Trends in neurosciences 36:152–162.

Stefan K, Kunesch E, Cohen LG, Benecke R, Classen J (2000) Induction of plasticity in the human motor cortex by paired associative stimulation. Brain 123:572–584.

Stefan K, Kunesch E, Benecke R, Cohen LG, Classen J (2002) Mechanisms of enhancement of human motor cortex excitability induced by interventional paired associative stimulation. The Journal of physiology 543:699–708.

Su F, Xu W (2020) Enhancing Brain Plasticity to Promote Stroke Recovery. 11.

Sui YF, Tong LQ, Zhang XY, Song ZH, Guo TC (2021) Effects of paired associated stimulation with different stimulation position on motor cortex excitability and upper limb motor function in patients with cerebral infarction. Journal of clinical neuroscience : official journal of the Neurosurgical Society of Australasia 90:363–369.

Tamura M, Spellman TJ, Rosen AM, Gogos JA, Gordon JA (2017) Hippocampal-prefrontal theta-gamma coupling during performance of a spatial working memory task. Nat Commun 8:2182.

Tarri M, Brihmat N, Gasq D, Lepage B, Loubinoux I, De Boissezon X, Marque P, Castel-Lacanal E (2018) Five-day course of paired associative stimulation fails to improve motor function in stroke patients. Annals of physical and rehabilitation medicine 61:78–84.

Ting WK, Huot-Lavoie M, Ethier C (2020) Paired Associative Stimulation Fails to Induce Plasticity in Freely Behaving Intact Rats. eNeuro 7.

Uy J, Ridding MC, Hillier S, Thompson PD, Miles TS (2003) Does induction of plastic change in motor cortex improve leg function after stroke? Neurology 61:982–984.

Volz MS, Finke C, Harms L, Jurek B, Paul F, Flöel A, Prüss H (2016) Altered paired associative stimulation-induced plasticity in NMDAR encephalitis. Annals of Clinical and Translational Neurology 3:101–113.

Wathugala M, Saldana D, Juliano JM, Chan J, Liew SL (2019) Mindfulness Meditation Effects on Poststroke Spasticity: A Feasibility Study. Journal of evidence-based integrative medicine 24:2515690×19855941.

Wessel MJ, Zimerman M, Hummel FC (2015) Non-invasive brain stimulation: an interventional tool for enhancing behavioral training after stroke. Frontiers in human neuroscience 9:265.

Wilke JT, Lansing RW, Rogers CA (1975) Entrainment of respiration to repetitive finger tapping. Physiological Psychology 3:345–349.

Yang CF, Feldman JL (2018) Efferent projections of excitatory and inhibitory preBötzinger Complex neurons. Journal of Comparative Neurology 526:1389–1402.

Zelano C, Jiang H, Zhou G, Arora N, Schuele S, Rosenow J, Gottfried JA (2016) Nasal Respiration Entrains Human Limbic Oscillations and Modulates Cognitive Function. The Journal of Neuroscience 36:12448–12467.

